# Exploring the influence of surface proteins on probiotic activity of *Lactobacillus pentosus* HC-2 in *Litopenaeus vannamei* midgut via Label-free quantitative proteomic analysis

**DOI:** 10.1101/546762

**Authors:** Yang Du, Han Fang, Mei Liu, Keyong Jiang, Mengqiang Wang, Baojie Wang, Lei Wang

## Abstract

Our previously work showed that *Lactobacillus pentosus* HC-2 as probiotic could improve the growth performance, immune response, gut bacteria diversity and disease resistance of *L*itopenaeus *vannamei*. However, the probiotic mechanism of was not fully characterized. In the present study, histology and proteomic analysis was performed to explore the influence of HC-2 surface protein on its probiotic effects to *L. vannamei* after feeding the intact surface proteins or the probiotic treated with lithium chloride (LiCl) to remove non-covalently bound surface proteins or no probiotic for four weeks. Histological observation found that feeding with normal HC-2 obviously improved the intestinal histology and enhanced the protective effect against pathogens damages, but fed with LiCl-treated HC-2 didn’t improve the intestinal environment. A total over 2,764 Peptides and 1,118 uniproteins were identified from *L. vannamei* midgut, 211 proteins were significant differentially expressed normal HC-2 group compared with control, 510 proteins were significant differentially expressed in LiCl-tread HC-2 group compared with control, and 458 proteins were significant differentially expressed in LiCl-tread HC-2 group compared with normal HC-2 group. GO/KEGG enrichment analysis of the significantly different proteins demonstrated that fed with normal HC-2 mainly induced immune response, metabolic, cell adhesion and cell-cell signaling related proteins up-regulation, which were contributed to bacteria adhesion and colonization in midgut to improve shrimp immune system and growth, but these proteins were suppressed after feeding with deprived surface proteins bacteria. Taken together, these results indicating that the surface proteins were indispensable for HC-2 to execute probiotic effects in midgut of shrimp.

## Introduction

*Litopenaeus vannamei*, is one of the most valuable crustacean aquaculture species worldwide because of its high nutrition value and tolerance to extensive salinity [1]. However, water environment deterioration, frequent disease outbreaks caused by viruses such as WSSV, YHV and IHHNV, or by bacteria like genus *Vibrio* are more prominent issues as a consequence of the rapidly growing shrimp aquaculture industry [2, 3]. Thus, there is international concern about dealing with the tough problem by supplying probiotic bacterial cells in food or in the aquatic environment to control the infectious diseases by strengthening the physique of aquatic animals [4, 5]. Among the available probiotics, lactic acid bacteria (LAB) are commonly used and advocated, such as *Lactobacillus pentosus, Lactobacillus helveticus, Lactobacillus delbrueckii, Lactobacillus acidophilus* and *Lactobacillusb plantarum* have been widely administered for the significant of improving host immune status, strengthening the host digestion, modulating the bacterial community, and antagonizing opportunistic pathogens [6-8].

The mechanisms of probiotic functionality LAB are not completely understood, but it is believed that the maximum probiotic effects can be achieved if the organisms adhere to mucus and/or intestinal epithelial cells [9]. It has recently suggested that surface proteins of lactobacilli bacteria participate in adhesion to epithelial cell lines, gastrointestinal mucins, or extracellular matrix proteins [10-12]. Indeed, except mediating binding ability, surface proteins are also involved in maintaining the shape of the bacteria, molecular sieve function, immunomodulation to the host, and providing extracellular enzyme binding sites [13-15]. Due to these proteins bind to the outermost layer of the bacteria with non-covalent bonds, make it possible that using denaturants lithium chloride (LiCl), guanidine hydrochloride (GuHCl), urea or metal chelating agents etc depolymerize them to monomer [16-18].

In previous work, we isolated a *Lactobacillus pentosus* HC-2 strain which has high antimicrobial activity against *Vibrio* pathogens and adhesive ability to intestinal mucosa, and regulated intestinal flora, and enhanced the growth performance, immune responses, and disease resistance after the *L. vannamei* fed with it [19-21]. The present study aim to further investigate the mediate function of surface proteins of *L. pentosus* HC-2 in the process of colonization and immune regulation of HC-2 to *L. vannamei*, label-free proteomic analysis was applied to characterize the proteins expression induced by surface proteins in midgut of shrimp fed with LiCl-treated HC-2.

## Materials and methods

### Bacterial growth and surface proteins shaving

*Lactobacillus pentosus* HC-2 (GenBank Accession No. KU995298) was previously isolated from the intestinal tract of fish (*Acanthogobius hasta*) by our laboratory [21], which was saved in −80 °C in de Man, Rogosa, and Sharpe (MRS) broth containing 20% (v/v) glycerol. After the recovery, the bacteria were cultured unstirred in MRS medium at 37 °C under anaerobic conditions.

Cell surface proteins shaving performed as previously [22]. Briefly, 500-mL culture of bacteria on the transition between late exponential and stationary phase (OD600 ≈1.7) were harvested centrifugation (3,000 × g, 10 min, 4 °C). Then, the cell pellets were washed three times with 1 M phosphate-buffered saline (PBS) containing 25% sucrose. After centrifugation, the bacteria cells were incubating the cells in 25ml of 5 M LiCl to stripe the surface associated proteins. After treatments, cells were collected and washed three times with autoclave sterilized seawater.

### Feeding trials

The experimental diets were prepared as previously that bacteria were resuspended in sterilized seawater and sprayed on basal commercial feed (containing crude protein 42%, crude fat 7%, ash 15%, and water 11%) at 5 × 10^8^ colony-forming units (CFU) g/feed [22]. A total of 600 shrimps (3.5 ± 0.06 g) were grown in twelve aquaria (60 L), each containing 50 shrimp. The experiments were designed as follows: C group, shrimp fed a basal commercial diet alone as the control; R group, shrimp fed a basal commercial diet + normal HC-2; L group, shrimp fed a basal commercial diet + LiCl treated HC-2. Three replicates were set in each feeding group. Keeping the fresh seawater (salinity, 30‰) at 30 ± 2 °C with continuous aeration and a 50% water change every day. Animals were fed three times per day, and the daily feeding rate was 10% of the body weight.

### Challenge test

After the feeding experiment, 25 shrimp were random selected from each aquarium and transferred to a tank with 30 L of seawater for challenge test. The live *Vibrio parahaemolyticus* E1 ATCC 17802 Strains was used for challenge, which was cultured aerobically in 2216E broth (Qingdao Hope Biol-Technology Co., Ltd) at 28 °C for 18 h. Preliminary experiment showed the appropriate bacteria dose was 10^7^ CFU/mL. During the challenge experiment, the shrimp were fed with basical diet.

### Histology of the Midgut

The histology determine was carried out as described in Sha et al. (2016) [20]. Five shrimps were freely selected from each treatment group upon termination of the feeding and challenge experiments and sampled the midguts to dissect and fix (60% absolute ethanol, 30% trichloromethane, 10% acetic acid) for 19 h. Following, the fixed tissues were dehydrated in ascending concentrations of alcohol (70, 80, 95, and 100%), cleared in toluene, embedded in paraffin, and sectioned at 10 μ m with a rotary microtome. The sectioned tissues were stained with hematoxylin and eosin, and images were obtained with a light microscope.

### Protein extraction and separation by 1D gel

Upon termination of the feeding experiment, the midguts of twenty shrimps from each treatment group were dissected and the intestinal contents was removed by flushing with sterile pre-cooled PBS. Total intestine proteins were extracted as the method described by Sengupta et al. (2011) [23] with some modifications. Pooled samples (1 g) were thoroughly grind into fine powder in liquid nitrogen with mortar and pestle and dissolved in 5 mL extraction buffer (0.5 M Tris-HCl (pH 7.5), 0.7 M sucrose, 0.1 M KCl, 50 mM EDTA, 40 mM DTT) at room temperature for 15min. After adding equal volume of Tris-phenol and shaking for 30 min, the upper phenolic phase was collected by centrifuging (8000 × g) for 5 min at 4 °C, and an equal volume of extraction buffer was added to the supernatant. For protein precipitation, four volumes of 0.1 M ammonium acetate in methanol were added and kept overnight at −20 °C. Protein pellet was collected after centrifugation at 8000 × g for10 min at 4°C, and washed thrice with ice-cold acetone at 4°C. The pellet was dried in vacuum for 2 h and then solubilized in 100 μL rehydration solution (8 M (w/v) urea, 0.1 M (w/v) Tris, 10 mM DTT). The concentration of protein was determined using the Bradford method [24]. Finally, proteins were loaded on 10% SDS-PAGE and separated at 120 V for 2 h, and visualized using colloidal Coomassie Blue after electrophoresis.

### Trypsin in-gel digestion

Protein gels were washed thrice with 50% acetonitrile (ACN)/50% NH_4_HCO_3_ (100mM) for10 min to destain, and dried in vacuum concentrator. The gels were dissolve in 200 μL 10 mM DTT containing 50 mM NH_4_HCO_3_ (pH 8.0) for 1h at 37 °C water bath, and then destained by 100 μL acetonitrile (ACN). Then, alkylation was performed with 55 mM iodoacetamide (Sigma-Aldrich)/50 mM NH4HCO3 (pH 8.0) for 30 min in the dark. The gel bands were alternately washed twice with 10 mM NH4HCO3 and 100% ACN, respectively. The gels were dried by Speed-Vac and digested with trypsin (0.01μg/μl) (Promega, Madison, WI) in 10 mM NH4HCO3 at 37 °C overnight. Digestion was stopped by adding 60% ACN/5% formic acid (FA) solution.

### Nanoflow liquid chromatography-tandem mass spectrometry

Prior to analyse the tryptic digest extracts using Thermo Scientic EASY-nLC 1000 System (Nano HPLC), the crude polypeptides were firstly desalted with a ChromXP Trap column (Nano LC TRAP Column, 3 μm C_18_-CL, 120 A, 350 μm×0.5mm, Foster City, CA, USA), and then eluted onto a second analytical column of Nano LC C_18_ reversed-phase column (3C_18_-CL, 75 μm × 15 cm, Foster City, CA, USA) under a linear gradient formed by mobile phases A (5% ACN and 0.1% FA) and B (95% CAN and 0.1% FA) at a flow rate of 300 nL/min for 120 min. Triple TOF 5600 MS (Foster City, CA, USA) was performed to automatically switch the TOF-MS and production acquisition in data-dependent mode by Analyst (R) Software (TF1.6).

### Protein identification

Three biological replicates were performed for the control, R and L group. The LC-MS/MS raw data were processed using the MaxQuant (version 1.5.2.8) for peptide/protein identification and quantification. MS/MS spectra was searched by the Andromeda search engine using a database consisting 28,384 sequences of the shrimp transcriptome, downloaded from NCBI. Search parameters were as follows: monoisotopic mass values; enzyme was trypsin; static Modification with C carboxyamidomethylation (57.021 Da); dynamic Modification was M Oxidation (15.995Da); Precursor ion mass tolerance ± 15 ppm; Fragment ion mass tolerance with ± 20 mmu; allowance of two missed cleavage site; false discovery rate (FDR) set as 0.01. Peptides identification with 95% confidence are considered “significant sequences”. For protein quantification, a minimum of two ratio counts was set to compare and normalize protein intensities across runs [25]. The absolute abundance of different proteins were then calculated using the intensity-based absolute quantification (iBAQ) algorithm, and iBAQ data were used for the t-test [26].

### Bioinformatics analysis

Gene ontology (GO) enrichment and Kyoto Encyclopedia of Genes and Genomes (KEGG) annotation for each protein in the search database were analyzed in GO (http://www.geneontology.org/) and KEGG Pathway database (http://www.genome.jp/Pathway), respectively. GO project provide three ontologies analysis, namely molecular functions (MF), cellular components (CC), and biological process (BP) [27]. Subcellular localisation for each protein was predicted according to GO annotation by Uniprot software (http://www.uniprot.org/). GO items without corresponding annotation were first deleted from the protein table, and then the IDs of listed proteins were plotted at the BP, CC, and MF levels. In addition, differentially expressed proteins (fold changes >1.5, p <0.05) were mapped to the GO database, and the number of proteins at each GO term was computed. The results from label-free proteomics were used as the target list. The background list was generated by downloading the GO database.

### Quantitative real-time PCR

RNA was prepared from midguts of ten *L. vannamei* that had been used for Real-time PCR analysis. Total RNA was extracted using the E.Z.N.A. HP Total RNA Kit (Omega Bio-Tek, Norcross, GA, USA) and reverse-transcripted into first-strand cDNA using the RevertAid First Strand cDNA Synthesis Kit (Thermo Fisher Scientific) according to the manufacturer’s instructions. Seven proteins including hemocyanin (Hem), C1q-binding protein (C1q), calreticulin (Cal), pyruvate kinase 2 (Pyr), integrin (Int), proliferating cell nuclear antigen (Pro), hemocyte transglutaminase (Htr) were selected to determine their mRNA levels, all of which were annotated to *L. vannamei* and were important proteins functioning in immune response, metoblic, adhesion and cell-cell signaling. The primer sequences were designed by Primer 5 (S1 Table). The reactions were carried out using Bio-Rad IQ™5 real-time PCR with a total volume of 20 m L (2*SYBR Green Mix (Vazyme Biotech): 10 m L, Primer: 1.0 m L, cDNA template: 5.0 m L, and PCR grade water: 4.0 m L). The qRT-PCR procedure was as follows: initial denaturation at 95 °C for 2 min; 40 cycles of amplification (95 °C for 15 s, 60 °C for 20 s, and 72 °C for 20 s). The cycle threshold (Ct) was measured, and the relative gene expression was calculated using the 2^-ΔΔCt^ method. *β*-actin gene was used as endogenous control. Three biological replicates and three technical replicates were done for all PCR experiments, and significance was determined at the P < 0.05.

## Results and discussion

### Histology of the Midgut

To investigate the effects of dietary Licl-treated probiotics on the midgut of the shrimp, a histological study was performed at the end of the feeding experiment and challenge assay as shown in Fig 1. Compared with the control group, the mucosae of R group improved more dense and more bloom (Fig 1B), but the mucosae of L group shrimps displayed to thin and loose (Fig 1C). After the shrimps were challenged by the *V. parahaemolyticus* E1, the mucosae shed and piled in the intestinal lumen, and the lamina propria exposed and appeared loose in the control and L groups (Figs 1D and 1F), especially, some individuals showed some reduced folding of the digestive epithelium in the crease. However, the shrimps fed with normal HC-2 appeared to no signs of necrotic enterocytes or cell damage (Fig 1E).

**Fig 1.**
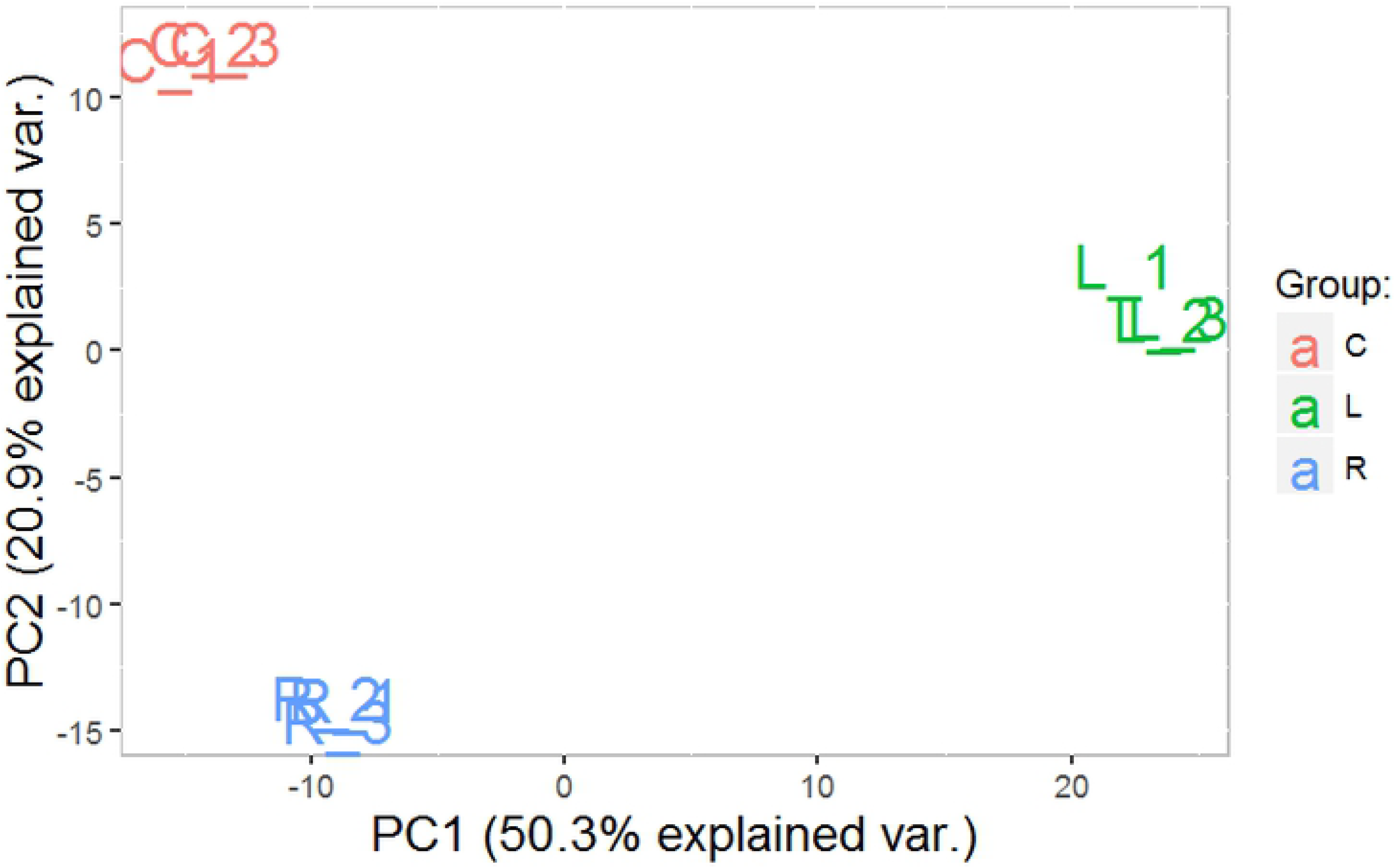
Histology with hematoxylin and eosin staining of the shrimp midguts after feeding with different diets for 4 weeks. Images A, B, C, D, E and F are arbitrarily chosen examples of the histology observed in three groups. A: Gut histology of shrimps were fed a basic diet; B: Gut histology of shrimps were fed a basic diet supplied with normal *L. pentosus* HC-2; C: Gut histology of shrimps were fed a basic diet supplied with LiCl-treated *L. pentosus* HC-2; D, E and F were showed the gut histology of shrimps in A, B and C respectively which were challenged by *Vibrio parahaemolyticus* E1. LP: lamina propria, M: mucosae, MV: microvilli, SCE: surface cell epithelium. Bar: 100 µm.

### Label-free proteomic analysis of intestine proteins of *L. vannamei*

In total, 2,810 proteins were detected. The differential protein expression among the three groups as shown in S2 Appendix, and proteins with fold change ≥ ± 1.5 and P < 0.05 were considered to be significantly differentially abundant. Pairwise comparison of intestinal protein with different levels among R/control, L/control, and L/R are illustrated in Fig 2, identified 210, 510 and 458 differentially abundant proteins, respectively. The numbers of upregulated proteins were 110, 134, and 85, respectively, whereas 101, 376, and 373 proteins, respectively, were down-regulated. The relationships among the experimental groups was performed by a PCoA analysis in proteins expression patter form. Samples of C, R and L group were clustered independently (Fig 3). To comprehensively analyze the impact of HC-2 and LiCl-treated HC-2 had on protein expression changes, the differentially abundant proteins were subjected to cluster analysis under different experimental conditions (Fig 4). Heat map showed that samples of C, R and L group were clustered respectively. The results of the heat map and PCoA were consistent to some extent, indicating that the proteins expression in C, R and L groups differed and that the proteins expression in the midgut was influenced by the addition of HC-2 and LiCl-treated HC-2.

**Fig 2.**
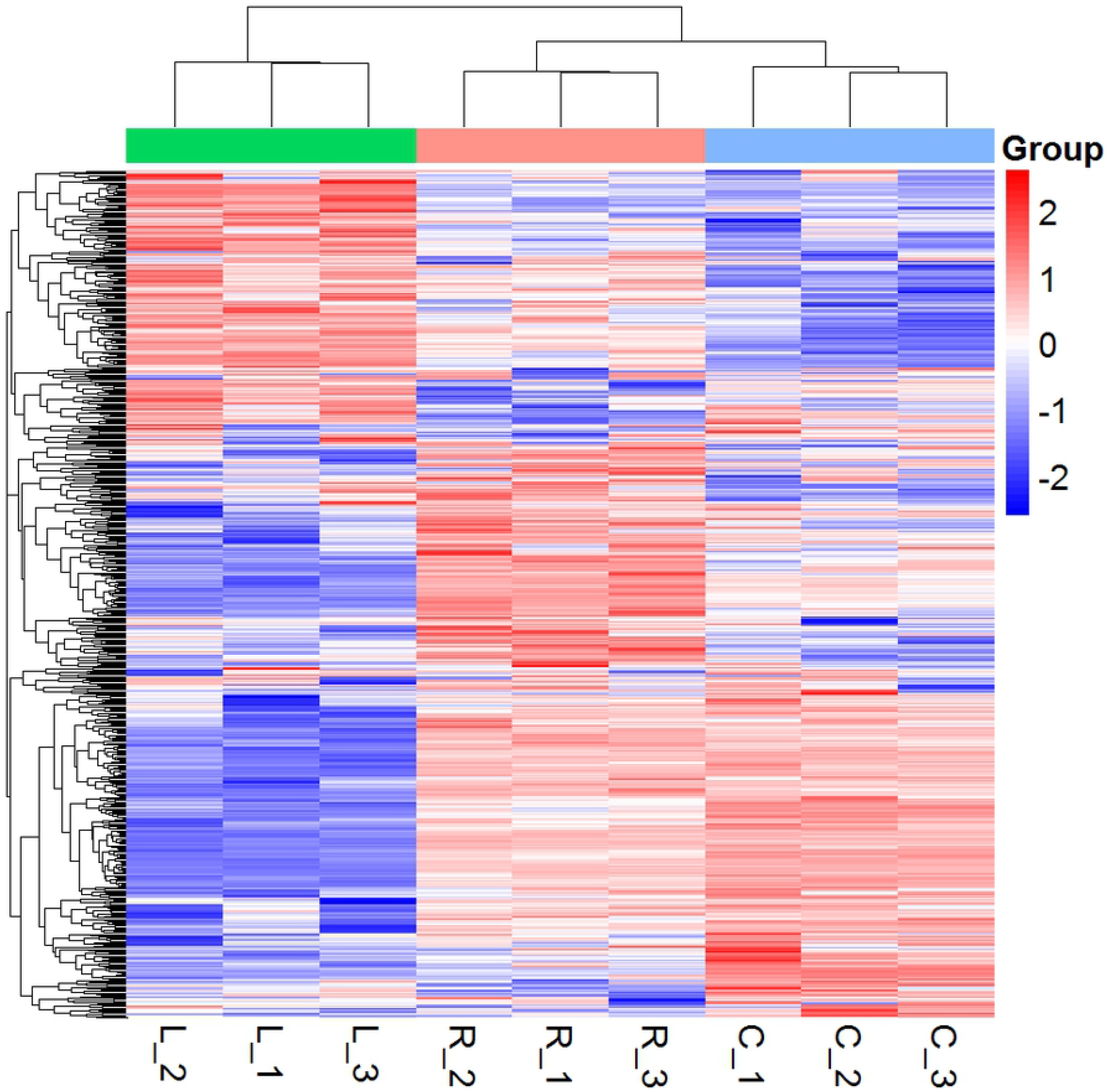
Volcano plot of changes in the levels of identified intestine proteins of shrimp analyzed using label-free quantitative proteomics after feeding with different diets. Note: C, shrimps were fed with basal diet; R, shrimps were fed with basal diet supplemented with normal *L. pentosus* HC-2; L, shrimps were fed with basal diet supplemented with LiCl-treated *L. pentosus* HC-2 (L).

**Fig 3.**
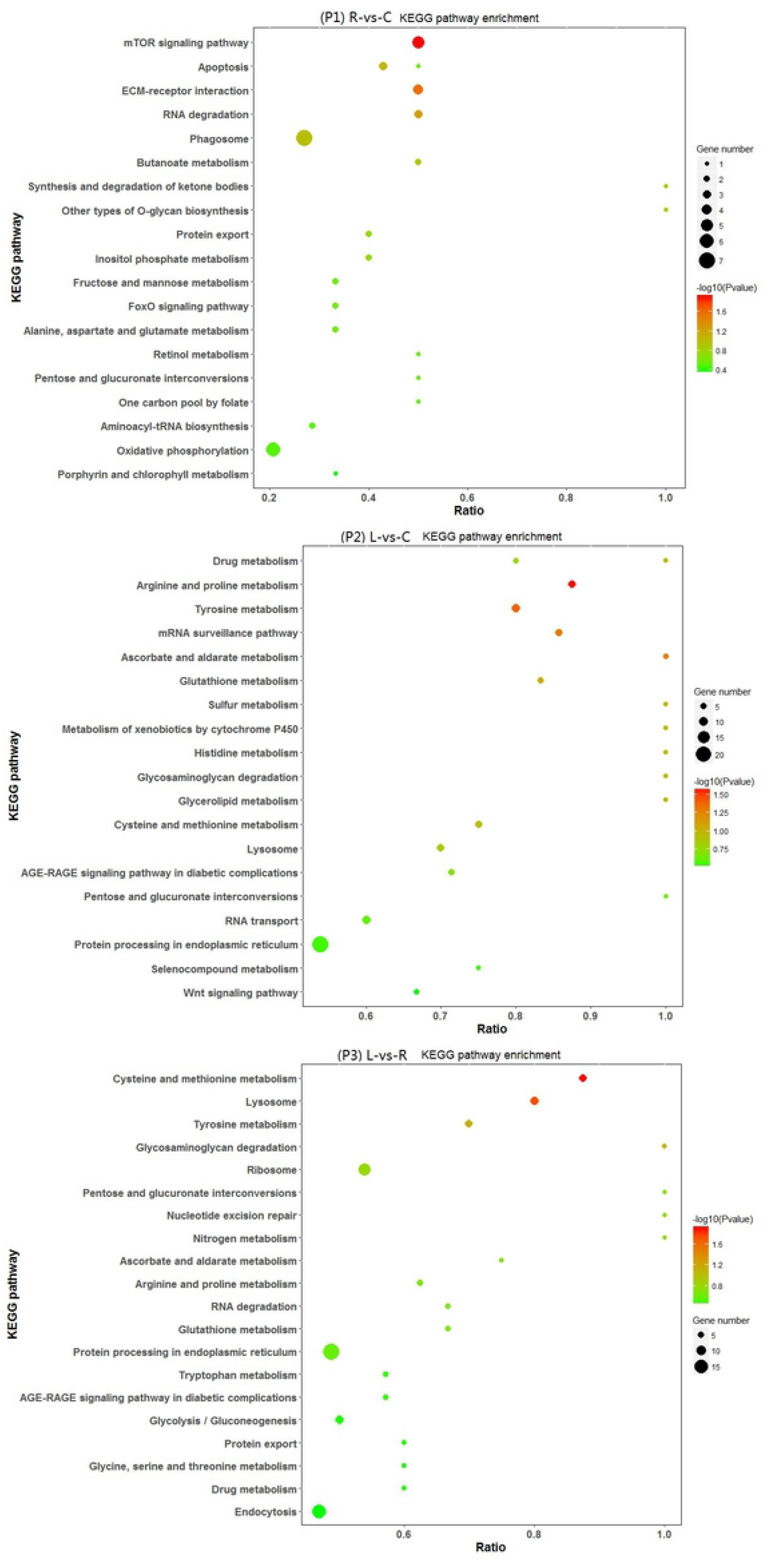
Principal coordinates analysis scores based on the Unifrac distance. PC1: the first principle component; PC2: the second principle component. Shrimps were fed a basal diet (C) or a basal diet supplemented with *L. pentosus* HC-2 (R), LiCl-treated *L. pentosus* HC-2 (L).

**Fig 4.**
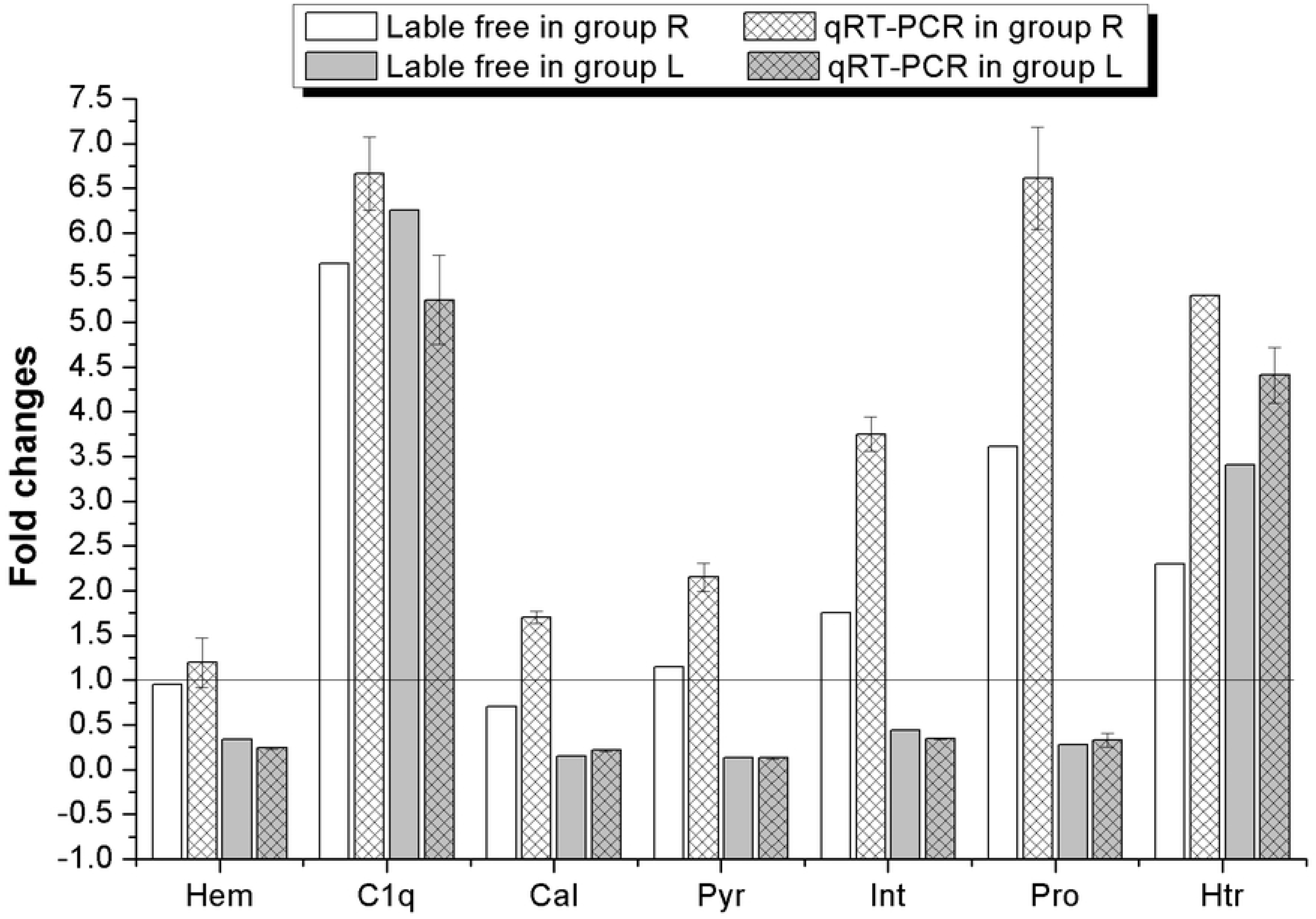
Heat map of the proteins expression diversity among the three groups (9 samples). Shrimps were fed a basal diet (C [C1, C2, C3]) or a basal diet supplemented with normal *L. pentosus* HC-2 (R [R1, R2, R3]), and LiCl-treated *L. pentosus* HC-2 (L [L1, L2, L3]).

### GO analysis of DEPs in *L. vannamei* midguts

Based on the gene ontology (GO) analysis in level 2 of biological process, cellular components and molecular functions associated with the significantly differentially abundant intestinal proteins (q-value < 0.05, and log2 |fold change| > 1.5) (Fig 5). Among the 210 differentially abundant proteins in the R/control comparison, 93 proteins played a role in 23 different biological processes, 139 proteins were related to cellular component and 46 proteins had distinct molecular functions. Compared with the control group, 510 differentially abundant proteins in the L group comprised 189 proteins that participated in 25 biological processes, 100 proteins had specific molecular functions, and 318 proteins were related to cellular components. Biological process analysis indicated that the transport, signal transduction, reproduction, immune system process, protein transport, transmembrane transport, embryo development, cell cycle, cell death, carbohydrate metabolic process, vesicle-mediated transport, growth, protein targeting, cell-cell signaling and cell adhesion processes involved the majority of proteins in R/control or L/control comparison. Cellular component analysis of R/control and L/control comparison revealed that the main differentially abundant proteins belonged to cytoplasm, membrane, nucleus, plasma membrane, mitochondrion, cytoskeleton, extracellular region and endoplasmic reticulum. Molecular function analysis revealed that most differentially abundant proteins were related to metal Ion binding and transmembrane transporter activity in both the R/control and L/control comparisons. Comparing L to R, 177 proteins played roles in the biological processes of transport, reproduction, signal transduction, vesicle-mediated transport, cell cycle, protein transport, immune system process, embryo development, growth, cell death, carbohydrate metabolic process, cell motility, cell-cell signaling, cell division, developmental maturation, protein targeting, membrane organization, cell adhesion; the cellular component of 290 differentially abundant proteins were cytoplasm, membrane, nucleus, mitochondrion, cytoskeleton, plasma membrane, extracellular region, endoplasmic reticulum, ribosome and Golgi apparatus; and 100 differentially abundant proteins in the molecular functions categories were related to metal ion binding, transmembrane transporter activity and signal transducer activity.

**Fig 5.**
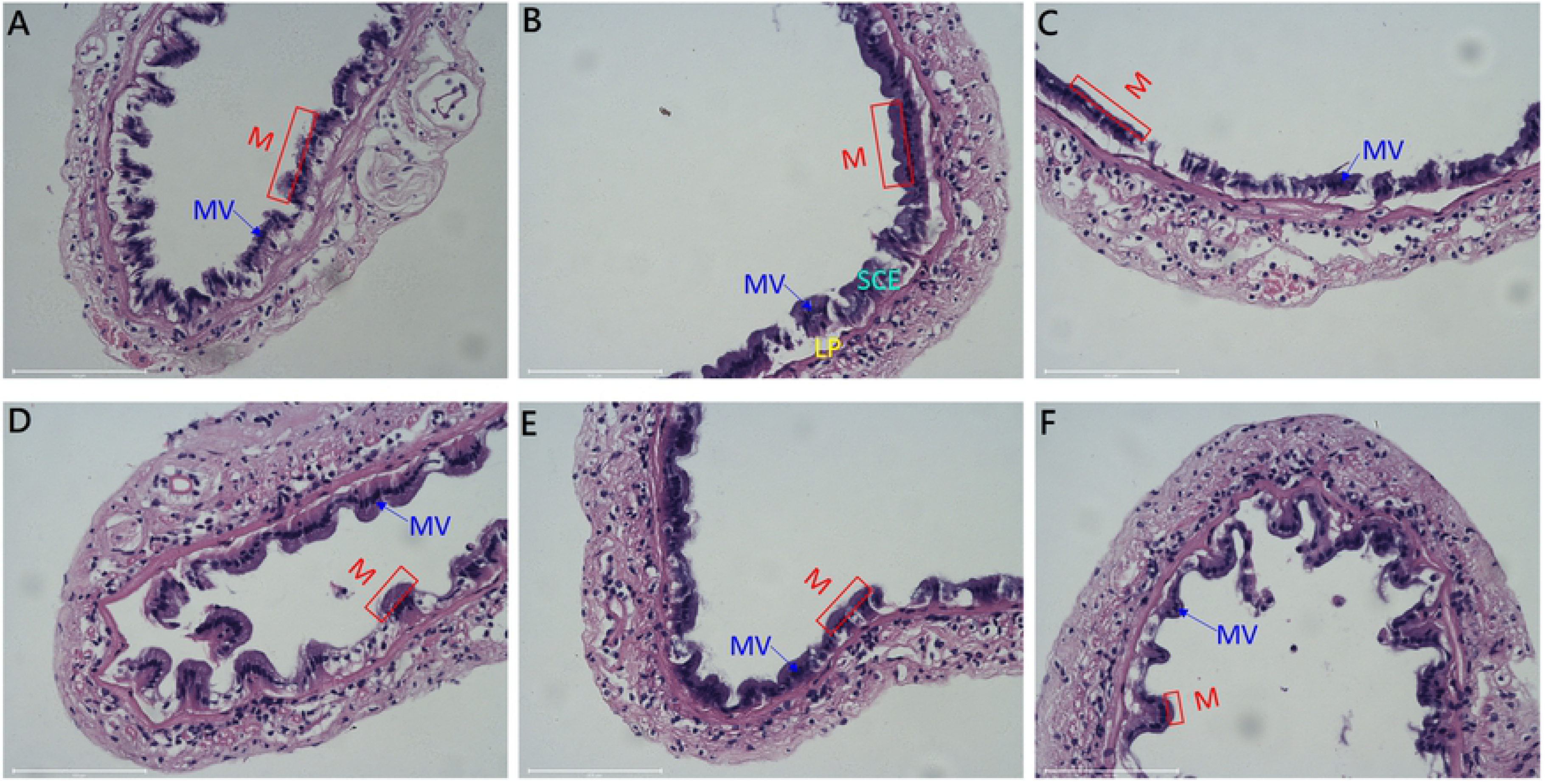
Functional categorization based on gene ontology (GO) in biological process, cellular components and molecular funcitons level analysis of significantly differentially abundant intestineal proteins. Shrimps were fed a basal diet (C) or a basal diet supplemented with L. pentosus HC-2 (R), LiCl-treated *L. pentosus* HC-2 (L).

### KEGG pathway analysis of the DEPs in *L. vannamei* midguts

KEGG pathway analysis was performed to determine the biological pathways that involved the differentially abundant proteins (q-value < 0.05, and log2 |fold change| > 1.5) induced by HC-2 and LiCl-treated HC-2 treatments fed in the diet (Fig 6). The DEGs between R group and the control group mainly enriched in mTOR signaling pathway, ECM-receptor interaction, RNA degradation, Apoptosis, Phagosome, Butanoate metabolism and Oxidative phosphorylation. The DEGs between the L and control group mainly enriched in Protein processing in endoplasmic reticulum, RNA transport, Tyrosine metabolism, Arginine and proline metabolism, Lysosome, mRNA surveillance pathway, Cysteine and methionine metabolism and Glutathione metabolism. The DEGs between R and L group mainly enriched in Protein processing in endoplasmic reticulum, Endocytosis, Ribosome, Lysosome, Glycolysis/Gluconeogenesis, Cysteine and methionine metabolism and Tyrosine metabolism.

**Fig 6.**
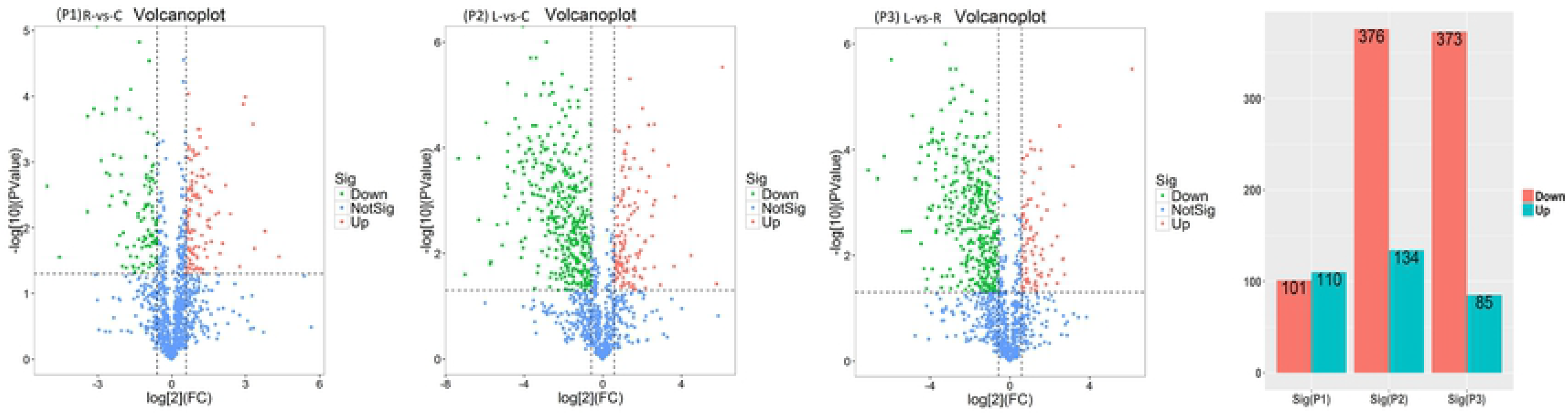
Distribution of differentially abundant proteins in shrimp midgut in KEGG pathways (Top 20). Note: Shrimps were fed a basal diet (C) or a basal diet supplemented with *L. pentosus* HC-2 (R), LiCl-treated *L. pentosus* HC-2 (L).

### Proteins potentially involved in shrimp immune response, metabolic, cell-adhesion and cell-signaling process

The GO enrichment analysis of biological process of the significantly different proteins involved in shrimp immune system process, cell-cell signaling process, cell adhesion and carbohydrate metabolic process among the three groups as shown in S3 Table 1. In the R/control group, 10 proteins involved in immune system process, 7 proteins involved in cell-cell signaling process, 5 proteins involved in cell adhesion and 4 proteins involved in carbohydrate metabolic process were significantly increased, and 3, 5 proteins involved in immune and carbohydrate metabolic process were significantly decreased. Among them, tyrosine-tRNA ligase, C1q-binding protein, tyrosine-protein phosphatase 69D-like and Neutral alpha-glucosidase AB were most up-regulated in the four process that the expression level reached 10.28, 5.66, 2.88 and 6.77 fold, respectively. However, in the L/control group, there didn’t induce more proteins up-regulation, and 22, 12, 6 and 18 proteins were participated in immune, cell-cell signaling, cell adhesion and carbohydrate metabolic process respectively were significantly down-regulated.

Based on the KEGG enrichment analysis, several of the proteins that were differentially expressed in shrimp fed with probiotic are involved in immune system process (mTOR signaling pathway, Apoptosis, Phagosome, Oxidative phosphorylation, MAPK signaling pathway, Lysosome, Protein processing in endoplasmic reticulum), Metabolism process (Arginine and proline metabolism, Tyrosine metabolism, Glutathione metabolism, Glycerolipid metabolism/Histidine metabolism, Cysteine and methionine metabolism, Fatty acid metabolism, Carbon metabolism), cell adhesion process (Focal adhesion, Tight junction, ECM-receptor interaction), and Cell signaling process (Calcium signaling pathway, Oxytocin signaling pathway, FoxO signaling pathway and Wnt signaling pathway) (S3 Table 2).

### Analysis of selected proteins affected by HC-2 and LiCl-treated HC-2 treatments

To validate the label-free based proteomic results, quantitative real-time PCR was used to analyze the transcripts of proteins found to be differentially abundant after HC-2 and LiCl-treated HC-2 treatments (Fig 7). The qPCR results showed that three proteins (Int, Pro and Htr) expressed higher than determined in R group proteome, the other proteins were consistent with the proteomics data, which further confirmed the reliability of label-free sequence.

**Fig 7.**
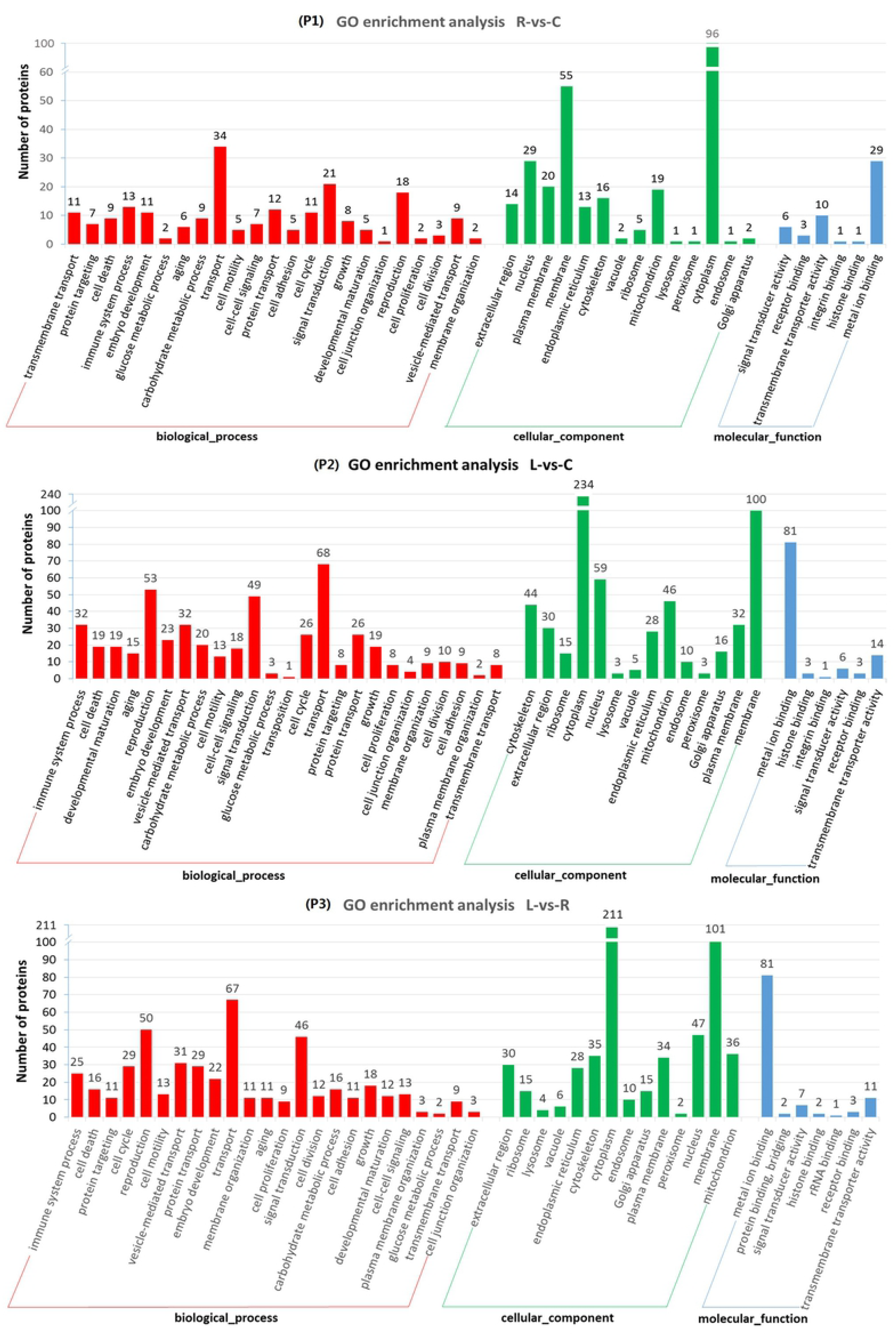
Validation analysis of label-free proteomics using quantitative real-time PCR to determine the selected proteins expression in midgut of *L. vannamei.* Note: R, shrimps were fed with basal diet supplemented with normal *L. pentosus* HC-2; L, shrimps were fed with basal diet supplemented with LiCl-treated *L. pentosus* HC-2 (L). Fold changes of proteins expression represent experimental group compared with control group.

## Discussion

Gastro-intestinal tract, the most important digestive and absorption organ in shrimps, where residing a large number of microorganisms with complex structures. These organisms depend on and restrict each other with hosts, and forming a unique intestinal micro-ecosystem in the long process of evolution [28]. Recent years, it is widely recognized that supplement with probiotics in aquaculture may stabilize the indigenous microflora, and normalize the host-microbe interaction, which is contribute to reduce the incidence of diseases [29]. Our previous work demonstrated that *L. pentosus* HC-2 has ideal probiotic effect to *L. vannamei*, but the probiotic action of surface components of HC-2 to shrimp is not clear. This work, to investigate the impact of surface proteins on probiotic effect of HC-2 to *L. vannamei*, proteomic analyses were conducted to using a label-free based LC-MS/MS approach to obtain protein data from three biological replicates.

Several studies demonstrated that dietary probiotic supplementation could improve the growth performance which was deemed to be attributed to intestinal physiology changes and gut epithelium morphology changes [30, 31], such as an improved intestinal microvillus structure and a greater absorptive surface area [32, 33]. In the present study, the changes in the intestinal microvilli and the folding of the digestive epithelium varied between dietary groups, and obvious improvement in intestinal histology was observed after shrimp fed with the normal probiotic HC-2, and the intestinal tissue was not damaged after the shrimp were challenged by *V. parahaemolyticus* E1. These results are similar to the findings of Merrifield et al. (2010) [33], who found that *Pediococcus acidilactici* - fed fish had significantly longer microvilli than other groups of fish, but are contrary to the findings of Sha et al. (2016) [20], who reported that dietary HC-2 didn’t improve the intestinal morphology of *L. vannamei*. These differently phenomena may be attributable to the bacteria concentration used in the dietary is too low (10^7^ CFU/g) than in this work (5×10^8^ CFU/g), which hinder the HC-2 to be the dominant microflora in the shrimp intestines to improve the intestinal morphology. However, no signs improvement in intestinal histology post the shrimp fed with LiCl-treated HC-2, instead, even to be more badly compared with the control shrimp that the mucosae showed to thin and loose after the shrimp challenged by pathogens. This results indicated that the surface proteins play important roles in probiotic function of HC-2 to improve the gut physiology and morphology.

With the intensive development of aquaculture and the frequent outbreaks of disease, varied probiotics have been developed to meet the demand of pollution-free immune enhancer. In shrimp farming, many authors have studied the influence of probiotic on the immune response. For example, Wang et al. (2010) [34] indicated that fed with *Lactobacillus* enhanced shrimp growth performance, increased digestive enzyme activities, and promoted non-specific immunity. Zheng et al. (2017) [35] also revealed that the administration of *Lactobacillus pentosus* AS13 effectively improved the shrimp growth performance, feed utilization, digestive enzymes and disease resistance. In present work, the proteomic analysis showed that fed with the normal HC-2 induced the proteins involved in immune system process (mTOR signaling pathway, Apoptosis, Phagosome, Oxidative phosphorylation, MAPK signaling pathway, Lysosome, Protein processing in endoplasmic reticulum) up-regulation, but many immune-related proteins were down-regulation in LiCl-treated HC-2 group shrimp midgut, which suggesting surface proteins play vital roles in mediation HC-2 enhance the shrimp intestinal immune response.

Several available genomic information descripted the metabolic activities of lactobacilli, which indicated that surface proteins are importance of carbohydrate metabolism in the host [36]. It has been reported that *Lactobacillus paracasei* or *Lactobacillus rhamnosus* probiotics supplementation of HBF mice exerted microbiome modification and resulted in altered hepatic lipid metabolism coupled with lowered plasma lipoprotein levels and apparent stimulated glycolysis, and also affected diverse range of metabolism pathways including amino-acid metabolism, methylamines and SCFAs [37]. In the present study, we found some proteins involved in carbohydrate metabolic process were significantly up-regulation after shrimp fed with normal HC-2, but proteins participated in other metabolic pathway including Arginine and proline, Tyrosine, Glutathione, Glycerolipid/Histidine, Cysteine and methionine, Fatty acid metabolism expressed insignificance. While, feeding with LiCl-treated HC-2 led to the overall downregulation of these metabolism related proteins, which indicated that surface proteins are importance in HC-2 regulation and maintenance of the shrimp intestinal metabolic.

Adhesion is the interaction of the bacteria surface structure (adhesin) attached with the surface receptors on the epithelial cells of the host, is the first step of bacterial colonization, and is the key for bacteria to grow, reproduce and functional exercise. Recent studies have indicated that the attachment of bacteria including the hydrophobicity and self-agglutination of the bacterial surface, lipoteichoicacid (LTA), exopolysaccharides (EPS) and related cell surface proteins to mucosal surfaces is the initial event in intestinal adhesion and colonization [38, 16]. Meanwhile, there are many surface proteins that mediate adhesion in lactobacillus have been reported, such as CmbA/Lar_0958, EF-Tu, GAPDH, GroEL, Lam29, MapA, MBF, Msa, Mub (Mub family), Pili, 32-Mmubp, FbpA and GroEL, etc [39-41]. Probiotics adhere to host intestinal mucus, intestinal epithelial cells, extracellular stroma by means of its surface proteins, and/or other bacteria lipodesmoic acid to effectively prevent pathogenic infection [42-44]. In present study, many related cell adhesion proteins were displayed significantly up-regulation in shrimp midgut after feeding with the normal HC-2, but the LiCl-treated HC-2 induced many cell adhesion proteins significantly down-regulation. Besides the adhesion ability, the surface proteins were studied have important functions in cell-cell signaling process, and interaction with the host immune system or environment [45]. In this study, we found that the proteins involved in cell-cell signaling pathway were up-regulated in shrimp midgut after fed with the normal HC-2, but the fed with surface proteins shaving bacteria induced these proteins decreased. These results indicated that surface proteins play crucial role in adhesion and colonization of HC-2 in the shrimp midgut, and were contributed to activation of a series of molecular signals communication with the surface cell of host.

In conclusion, fed with normal HC-2 obviously improved the intestinal histology and enhanced the protective effect against pathogens damages, but fed with LiCl-treated HC-2 didn’t improve the intestinal structure. GO and KEGG enrichment analysis of significantly proteins in R/control and L/control indicated that most proteins were involved in immune system process, metabolic process, adhesion process, and cell-cell signaling process. However, these proteins were significantly up-regulation in shrimp midgut after feeding the normal HC-2, and were significantly down-regulation in shrimp fed with LiCl-treated HC-2. The results in present work indicated that surface proteins play an important roles in mediation of HC-2 to improve intestinal histology, immune response, metabolic, adhesion and signaling communication in midgut of shrimp, which might provide a base data to understand the probiotic mechanism excised by HC-2.

## Supporting information

S1 Appendix. Primers. (PDF)

S2 Appendix. Different expression proteins (Excel)

S3 Appendix. Significantly different expression proteins (PDF)

## Acknowledgments

We thank Dr Wenbin Zhan (Laboratory of Pathology and Immunology of Aquatic Animals, Ocean University of China), for providing *V. parahaemolyticus* E1, and we are grateful to all the laboratory members for their technical advice and helpful suggestions.

